# The ECF sigma factor PvdS regulates the type I-F CRISPR-Cas system in *Pseudomonas aeruginosa*

**DOI:** 10.1101/2020.01.31.929752

**Authors:** Stephen Dela Ahator, Wang Jianhe, Lian-Hui Zhang

## Abstract

During infection, successful colonization of bacteria requires a fine-tuned supply of iron acquired via iron transport systems. However, the transport systems serve as phage attachment sites and entry portals for foreign nucleic acid. Most bacteria possess the CRISPR-Cas system, which targets and destroys foreign nucleic acids and prevents deleterious effects of horizontal gene transfer. To understand the regulation of the CRISPR-Cas system, we performed genome-wide random transposon mutagenesis which led to the identification of the Extracytoplasmic Function (ECF) Sigma factor, PvdS as a regulator of the Type I-F CRISPR-Cas system in *P. aeruginosa*. We show that under iron-depleted conditions PvdS induces the expression of the type I-F CRISPR-Cas system. This regulatory mechanism involves direct interaction of PvdS with specific binding sites in the promoter region of *cas1*. Furthermore, activation of the CRISPR-Cas system under iron-depleted conditions increases horizontal gene transfer (HGT) interference and adaptation. The PvdS activation of the CRISPR-Cas system under iron limitation highlights the versatility of the *P. aeruginosa* in multitasking its regulatory machinery to integrate multiple stress factors.

**Importance:** *P. aeruginosa* infects a wide range of host organisms and adapts to various environmental stress factors such as iron limitation due to its elaborate regulatory system. *P aeruginosa* possesses the type I-F CRISPR-Cas system as a defense mechanism against phages infection and HGT. This work highlights the ability of *P. aeruginosa* to multitask its iron regulatory system to control the CRISPR-Cas system under a physiologically relevant stress factor such as iron limitation where the bacteria are vulnerable to phage infection. It also adds to the knowledge of the regulation of the CRISPR-Cas system in bacteria and presents a possible target that could prevent the emergence of phage resistance via the CRISPR-Cas system during the development of phage therapy.

## Introduction

The adaptation of *Pseudomonas aeruginosa* to stress conditions, effective sensing and response to varying environmental factors are due to its versatile and extensively interconnected regulatory systems. These regulatory systems enable the bacteria to alter its regulatory pathways to release factors that will enable it thrive under different stress conditions and colonize biotic and abiotic surfaces. As an opportunistic pathogen, *P. aeruginosa* can colonize and infect various parts of the human body with a breach of the immune barrier. One major factor required for its successful colonization during infection is iron (1). Iron remains a universal requirement for bacterial growth therefore multiple systems are dedicated to its acquisition from the host and its environment. In human hosts, iron is usually bound to heme, myoglobin, transferrin and lactoferrin whiles it is maintained in very small amounts as a free soluble form in the environment (1). Thus, the problem with iron is not one of abundance, but of availability in aerobic environments. Hence to acquire iron bacteria produce high-affinity iron chelators called siderophores and express transport systems for their uptake (2).

Under iron-limited conditions, *P. aeruginosa* produces two major siderophores, pyoverdine and pyochelin which are transported via the TonB-dependent FpvA and FptA respectively (3)(4). Production of siderophores and expression of their transporters are active processes which can be metabolically costly to the bacteria. Hence, they are correspondingly upregulated by the positive regulator, Extracytoplasmic function (ECF) sigma factor, PvdS under iron-depleted conditions and downregulated by the Ferric Uptake Regulator (Fur) under iron-rich conditions (5). Additionally, transport of the siderophores via the uptake systems are dependent on the proton motive force generated by TonB protein. The expression of the TonB-dependent transporters is induced by iron limitation (6). *P. aeruginosa* possesses three TonB proteins (TonB, TonB2, and TonB3) and a host of TonB-dependent siderophore transport systems which show varying degrees of functions in iron transport (7)(8)(9).

A trade-off in expressing TonB-dependent iron transporters is the predisposition to bacteriophage infection as they serve as attachment sites and entry portals for phage nucleic acids (10)(11)(12). Additionally, the TonB-dependent electromotive force required for siderophore-iron transport enables the transfer of phage nucleic acid into cells during the phage infection process (13)(14). This makes *tonB* mutants resistant to phage infection (12). As phages outnumber bacteria in nature, bacteria therefore face a phage infection challenge in their quest to acquire iron for their growth and colonization.

As a defense mechanism, bacteria have evolved the inducible immune defense mechanism, the Clustered Regularly Interspaced Short Palindromic Sequences (CRISPR)-CRISPR associated (Cas) Protein system, which targets and destroys foreign nucleic acid from phage infection and HGT. Some strains of *P. aeruginosa* contain the Type I-F CRISPR-Cas system which consists of 6 *cas* genes flanked by two CRISPR arrays. The CRISPR-Cas provides adaptive immunity against new and previously encountered bacteriophages and plasmids (15). Following entry of foreign nucleic acids, the CRISPR-Cas system incorporates short segments of the nucleic acid into the CRISPR array which act as immunological memory in a process known as adaptation (16). The CRISPR arrays are transcribed and processed into crRNAs which guide the Cas proteins to complementary sequences of foreign nucleic acid. The crRNA guided Cas proteins complex then cleaves and destroys the foreign nucleic acid in a process termed interference (17)(18)(19).

As an inducible system, it is strictly regulated to avoid the metabolic burden due to its constitutive expression and autoimmunity (20)(21, 22). The CRISPR-Cas systems in bacteria are regulated in response to population dynamics and environmental factors such as Quorum Sensing, glucose availability, temperature(23)(21)(15)(24)(25). These factors are known to predispose the bacteria to phage infection and also influence their ability to defend against phage infection via the CRISPR-Cas system.

Bacteria encounter various biotic and abiotic stress factors in the external environment and host that make them vulnerable to phage infection. Here we show an iron-dependent regulation of the type I-F CRISPR-Cas system via the ECF sigma factor, PvdS in *P. aeruginosa*. This regulatory pathway of the CRISPR-Cas system by PvdS in response to iron limitation highlights the ability of *P. aeruginosa* to adapt by evolving contingency pathways under conditions when it is most vulnerable to phage infection.

## Results

### Identification of PvdS as an iron-dependent regulator of the Type I-F CRISPR Cas system

We used the *P. aeruginosa* UCBPP-PA14 which contains the type I-F CRISPR-Cas system with two CRISPR loci flanking 6 *cas* genes (*cas1, cas3, csy1-csy4*)(Fig. 1A). To identify the regulators for the expression of Type 1-F CRISPR-Cas system, we performed a genome-wide transposon mutagenesis with the mariner transposon in the wild type PA14 bearing a *cas1* promoter-*lacZ* fusion construct (PmeP*cas1-lacZ*) (Fig. 1A). Using this construct, transposon mutants with reduced *cas1* expression were visually inspected and selected for β-galactosidase analysis. White colonies were due to transposons inserted in the construct and were not selected for further analysis. Following the visual inspection, we identified several transposon mutants exhibiting less blue coloration compared to the wild type. Transposon insertion site mapping revealed several genes, with a notable one encoding the ECF sigma factor, PvdS located upstream of the CRISPR-Cas genes (Fig. 1A). Since PvdS is activated under iron-depleted conditions, we adapted the iron-depleted media, LB + 2,2-dipyridyl (LB+2,2DP) (26)(8) to study its regulatory effect on the CRISPR-Cas system. Growth analysis of the strains in LB and LB+2,2DP showed that the latter supports the growth of *pvdS* mutants (Fig. S1), therefore this media was used for our downstream analysis. Despite the minor growth defect of *pvdS* mutant under iron-depleted conditions compared to the wild type (Fig. S1), it was not sufficient to influence the expression of the CRISPR-Cas system. In the wild type background expression of the *cas1* was upregulated under iron-depleted conditions compared to the normal conditions (LB) (Fig. 1B). Furthermore, expression of the *cas*1 was significantly reduced in the *pvdS* mutant whereas deletion of *pvdS* did not effect *cas* gene expression under normal conditions (Fig. 1B). Additionally, the expression of the *cas* gene in the *pvdS* mutant was rescued when we complemented *pvdS* in trans under iron-depleted conditions (Fig. S1B). In general, the expression of the *cas* genes was upregulated under iron-depleted conditions compared to normal condition (fig S1C).

**Figure 1:**
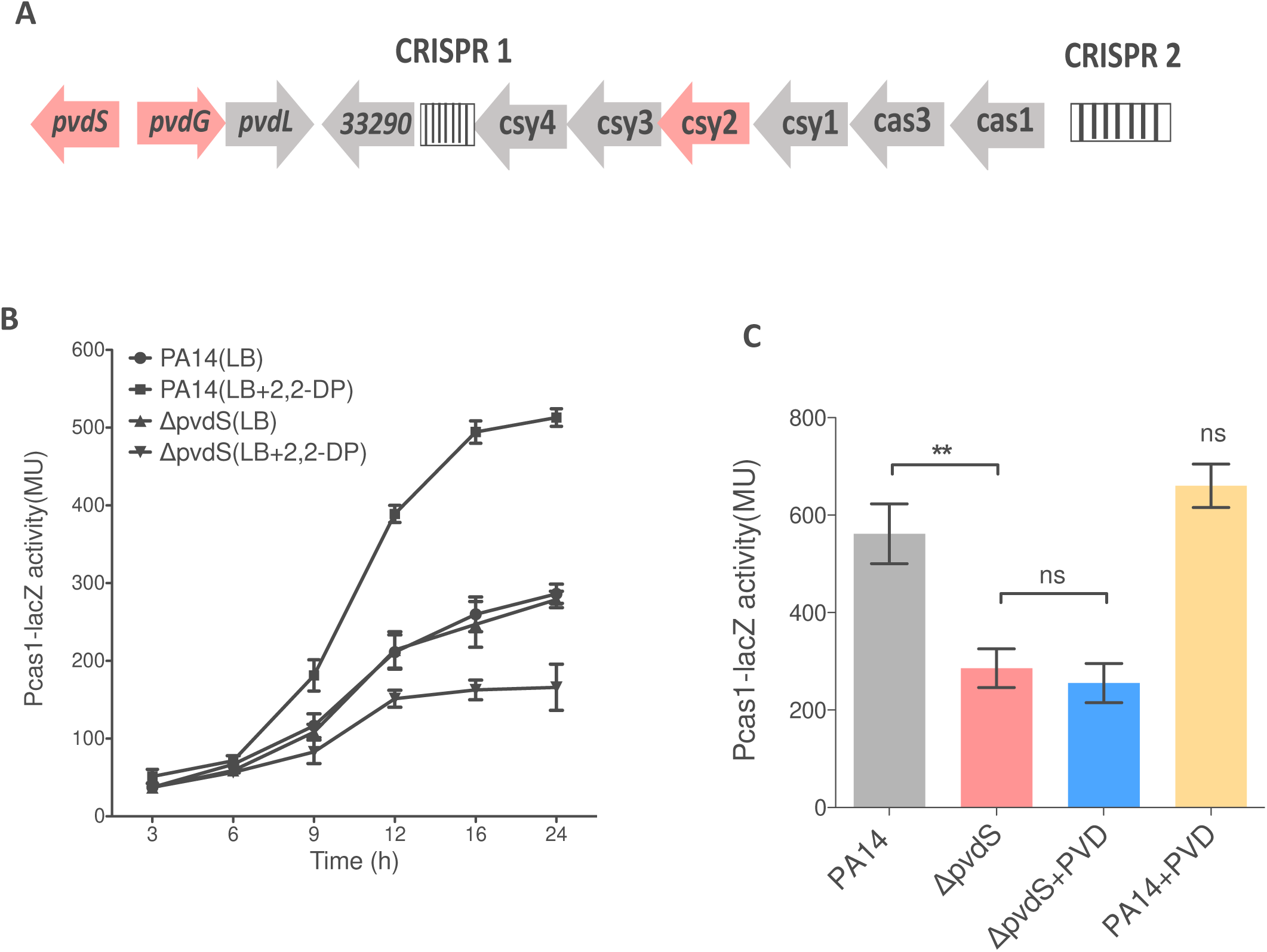
Identification of PvdS as a regulator of type I-F CRISPR-Cas system. **A**. Genome map of *pvdS* and the CRISPR-Cas system in *P. aeruginosa* UCBPP_PA14. **B**. Time course β-galactosidase assay for the *Pcas1-lacZ* activity in Wild type (PA14) and in-frame deletion mutants of *pvdS* grown in LB and LB+2,2-DP.The Pcas1-lacZ activity was normalized to the OD_600_ at each time point. Data represent the mean ± SD of n=3 independent experiments. **C**. Expression of *cas1* in wild type and *pvdS* mutants grown in LB+2,2-DP with or without exogenous pyoverdine (PVD), 30µM. Data represent the mean and SD of n=3 experiments. Statistical analysis was performed using student t-test (**P<0.001, ns=non-significant).

The expression of *pvdS* is repressed by Fur when iron is abundant in the environment thus, resulting in the downregulation of the PvdS-regulated genes. However, under iron-depleted conditions, Fur relieves PvdS, allowing it to bind to the consensus binding sites in the promoter region of genes and inducing their expression (26). To investigate whether Fur influences the PvdS-CRISPR-Cas regulation, we analyzed the expression of the *cas* genes in the *fur* and *pvdS* mutants under iron-depleted and normal conditions. In the *fur* mutant expression of the *cas* genes were upregulated under both normal and iron-depleted conditions relative to the wild type (Fig. S2A and S2B). This was contrary to the expression of *cas* genes in the *pvdS* mutants as they were downregulated under iron-depleted condition (Fig S2A). This shows that the deletion of *fur* derepressed the *pvdS* even under normal conditions and therefore allows PvdS to regulate the *cas* genes. This is in agreement with the Fur-PvdS regulatory system shown by previous studies (5)(27)(28). From our analysis, exogenous pyoverdine (PVD) did not significantly increase *cas* gene expression in both wild type and *pvdS* mutant (Fig. 1C), which signifies the possibility that PvdS regulation of the CRISPR-Cas system may be independent of siderophore production and uptake.

### PvdS binding site in *cas1* promoter is required for CRISPR-Cas regulation

PvdS directly regulates its target genes by interacting with a consensus binding site in their promoter region. This interaction and regulation are facilitated by the formation of a complex with Core RNA polymerase (cRNAP)(26). The examination of the *cas1* promoter region revealed the presence of two PvdS binding sites (Fig 2A). To investigate whether the predicted PvdS binding site are required for interaction and regulation of the *cas* genes, we created an integrative *cas1* promoter-*lacZ* reporter strain with altered PvdS binding site sequence and examined the effect on *cas1* expression under iron-depleted conditions (Fig 2B). Initially, we investigated the effect of the mutations on the interaction between PvdS and PvdS-cRNAP complex with the *cas1* promoter region. The binding assay revealed that the mutation of one binding site did not affect the interaction between PvdS-cRNAP and the *cas1* promoter region (fig 2C). However, PvdS-cRNAP binding was abolished when the sequences of both binding sites were altered (fig 2C). Also, no interaction was observed between the intact and altered PvdS binding sites in the absence of cRNAP (Fig. S3), which depicts a cRNAP-dependent PvdS interaction.

**Figure 2.**
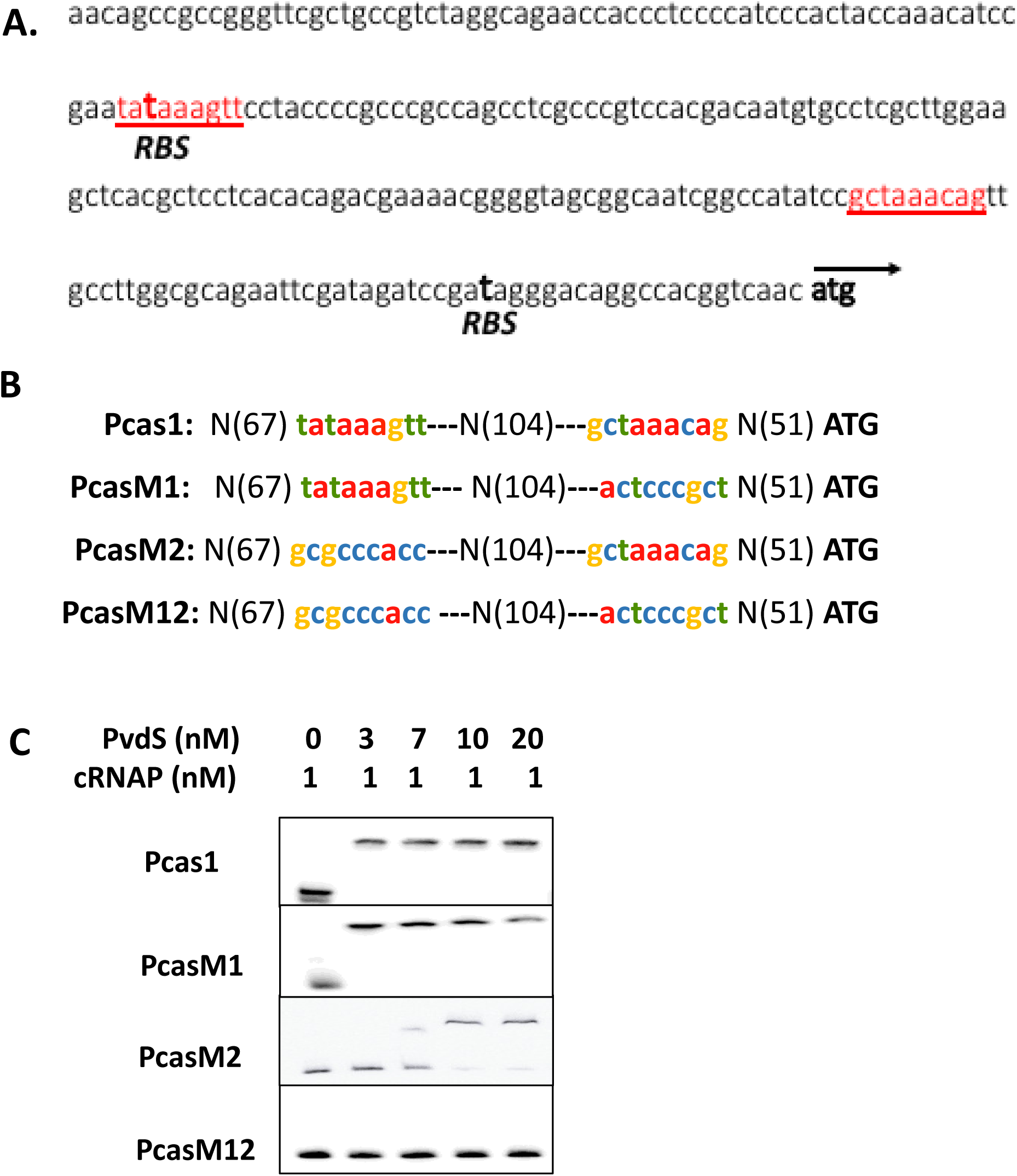
PvdS interacts with the *cas1* promoter. **A**. The DNA sequence of cas1 promoter region. The underlined sequence represents the PvdS binding sites. The ribosomal binding sites are in bold and labeled as RBS. **B**. The two intact PvdS binding sites in *P. aeruginosa cas1* promoter is labeled as Pcas1. PcasM1 and PcasM2 are the promoter sequences with a single altered PvdS binding site separated by 104 nucleotide N(104). PcasM12 contains both altered PvdS binding sites at M1 and M2. **C**. EMSA analysis of PvdS-cRNAP interaction with biotin-end labeled probes of Pcas1, PcasM1, PcasM2, and PcasM12.

Furthermore, alteration of either PvdS binding sites significantly reduced the expression of *cas1* in the wild type PA14 under iron-depleted conditions (Fig3A). The decrease in *cas1* expression was more significant when we altered both binding sites in the wild type PA14 compared to the single binding alteration in the wild type background (Fig 3A). Additionally, alteration of single PvdS binding sites resulted in similar expression as the intact sequence in the *pvdS* mutant background (Fig. 3). These results show that PvdS interaction with a consensus binding site in the promoter region of *cas1* is required for *cas* gene expression. Since PvdS is known to require multiple sites for full activity *in vivo* and *in vitro* (26), the decrease in *cas* gene expression as a result of mutations in the binding sites may imply the requirement of both sites for optimal regulation by PvdS.

**Figure 3.**
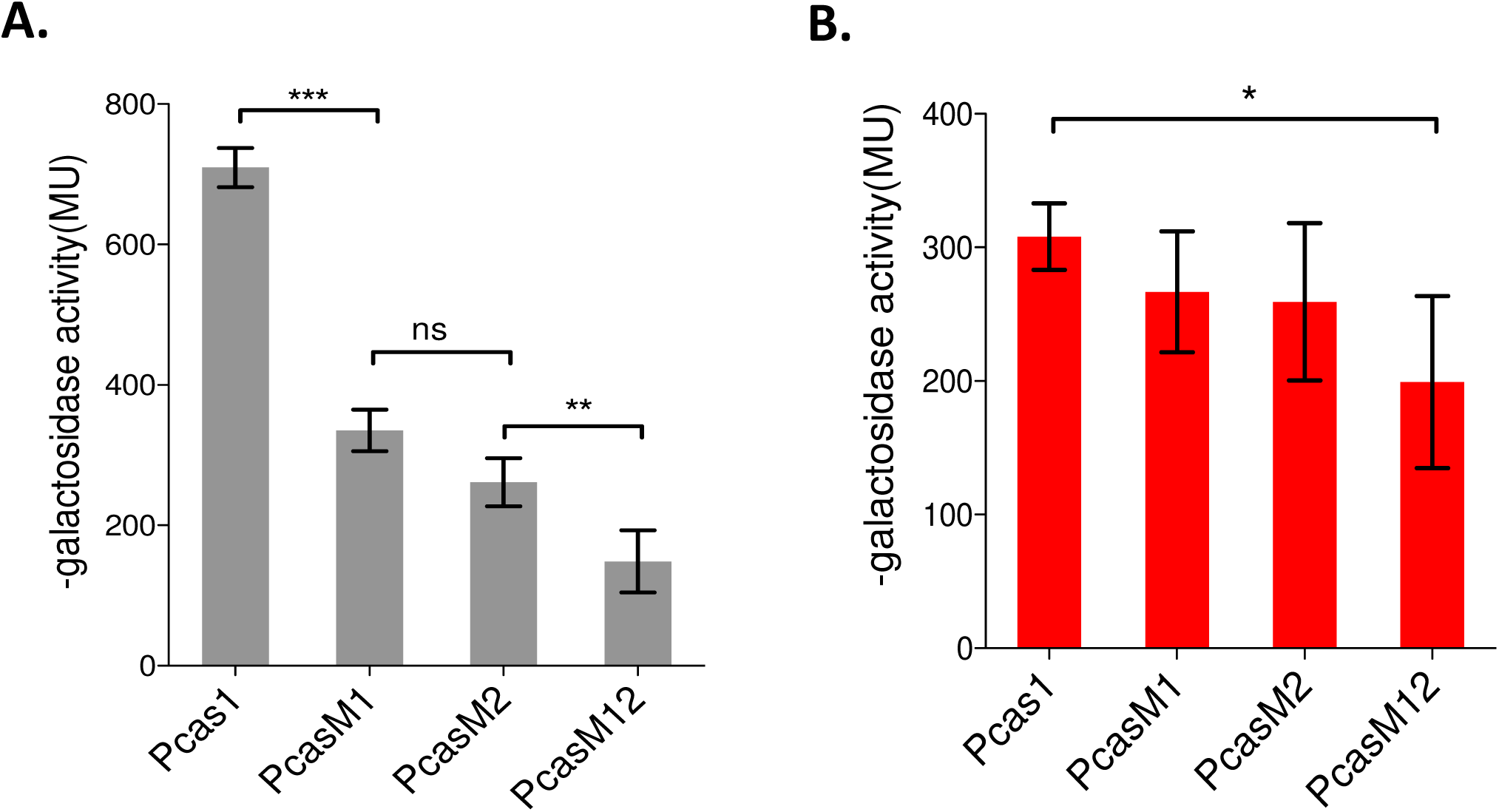
PvdS binding site in *cas1* promoter is required for activation. **A** and **B**. β-galactosidase activity of Pcas1-lacZ, PcasM1-lacZ,PcasM2-lacZ and PcasM12-lacZ in PA14 (**A**) and *pvdS* mutant(**B**) backgrounds measured after 16 hours post inoculation. Data represents mean± SD of n=3 independent experiments. Statistical analysis was performed using the student t-test (***P=0.0001, **P=0.002,*P=0.01 ns=non-significant P=0.08)

### PvdS influences CRISPR-Cas interference of horizontal gene transfer and adaptation

Since PvdS regulates the *cas* genes expression under iron-depleted condition, we further investigated how this regulation influences the function of the CRISPR-Cas system in interfering with HGT and the acquisition of spacers from the foreign nucleic acid. Acquisition of spacers is required for the development of immunological memory against specific nucleic acid sequences (29). To test the impact of PvdS on the interference and adaptation of the CRISPR-Cas system, we engineered CRISPR-targeted constructs containing sequences similar to CRISPR 2 spacer 1 of the type I-F CRISPR-Cas system in PA14. In the three constructs designed, pUCPTsp1 contained matching sequences to CRISPR 2 spacer while pUCPTSp2n and pUCPTSp4n contained 2 and 4 nucleotide substitutions respectively after the GG PAM sequence recognized the Type I-F CRISPR-Cas system. These mutations in the protospacer reduce interference but can initiate primed adaptation (21)(15). The naïve plasmid used lacked the protospacer and the consensus GG PAM sequence.

We initially quantified the retention of the naïve and the CRISPR-targeted (pUCPTSp1) plasmid transferred into the *pvdS* mutant and the wild type PA14 under iron-depleted and normal conditions (fig 4). Retention of the CRISPR-targeted plasmid was significantly reduced in the wild type compared to the *pvdS* mutant when passaged over five days in the iron-depleted media (Fig. 4A). As expected, there was no significant loss of the naïve plasmid when transferred in both the wild type and *pvdS* mutant after passage over five days in both normal conditions and iron-limited conditions (Fig. S5A and S5B). However, under normal conditions, the wild type and *pvdS* mutant exhibited similar levels of CRISPR-Cas targeted-plasmid loss (Fig. 4B). These results show that PvdS is required for the induction of the CRISPR-Cas plasmid interference under iron-depleted conditions.

**Figure 4.**
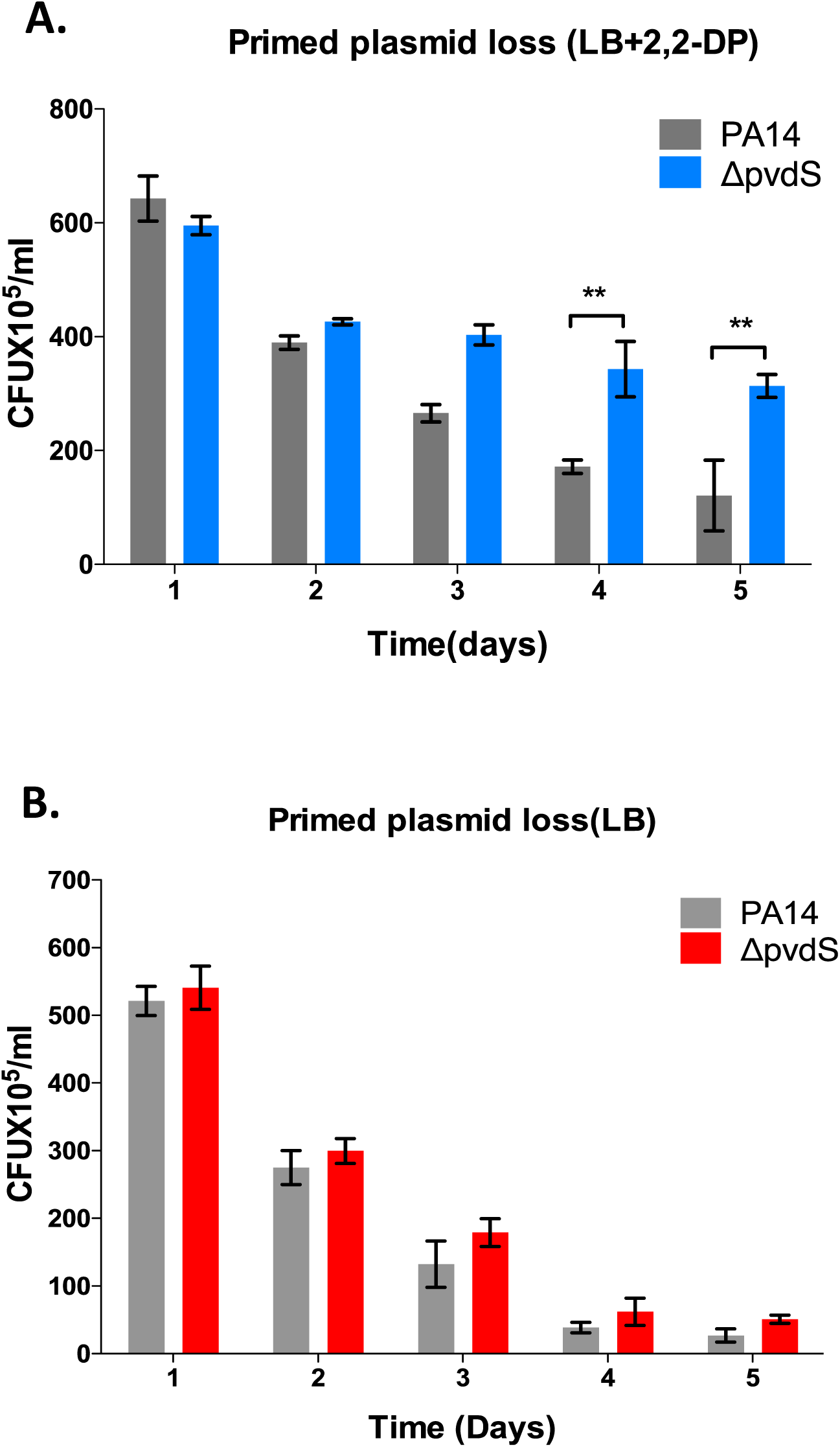
PvdS regulates CRISPR-Cas interference. **A** and **B**. Retention of CRISPR-Cas targeted plasmid in PA14 and *pvdS* mutant strains after 5 days passage in LB+2,2-DP (**A**)and LB (**B**). Plasmid loss was scored by counting positive colonies on plates LB plates containing carbenicillin. Data represent the mean ± SD (*n* = 3). Statistical significance was calculated using student t-test(**P≤ 0.001).

Additionally, we investigated the influence of PvdS on CRISPR-Cas spacer acquisition which is depicted by an expansion of the CRISPR2 array. We targeted the CRISPR2 which has been previously shown to have a higher adaptation frequency in *P. aeruginosa* (20)(15). The *P. aeruginosa* wild type PA14, *pvdS and cas3* mutants containing CRISPR-Cas targeted plasmids (pUCPTSp1, pUCPTSp2n, and pUCPTSp4n) were passaged for five days in both iron-depleted and normal media. Three successive adaptation assays were performed in the strains to test the versatility of the *P. aeruginosa* CRISPR-Cas system to incorporate multiple spacers into its array. As expected, deletion of *cas3* resulted in no expansion of the CRISPR2 array under both iron-depleted and normal conditions compared to the wild type (Fig. 5A). Also, a significantly lower expansion (∼60% less than wild type) of the CRISPR2 array was detected in the *pvdS* mutants compared to the wild type under iron-depleted conditions. However, there were no significant differences in spacer acquisition observed under normal conditions between the wild type and the *pvdS* mutant (Fig. 5A).

**Figure 5.**
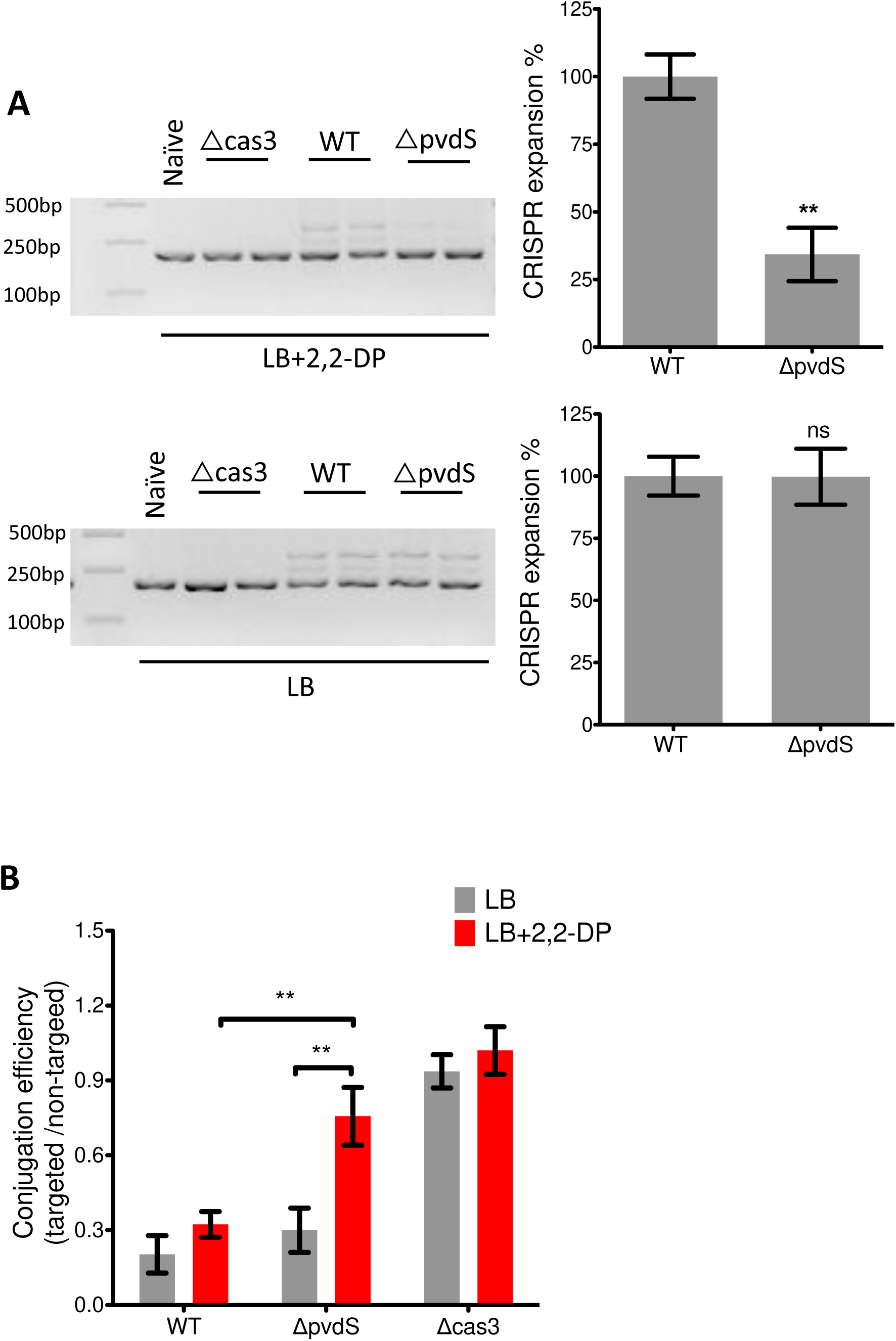
PvdS influences CRISPR-Cas regulation of HGT and spacer acquisition. **A**. Gel electrophoresis showing acquisition of spacers in the CRISPR2 locus of wild type PA14, *pvdS*, and *cas3* mutants strains passaged in LB and LB+2,2-DP for 5 days. As a negative control, wild type was transformed with the non-targeted plasmid (naïve). The adaptation events are signified by the expansion of the arrays and 60bp extension of the CRISPR locus. The image is a representative of 4 similar replicates. Quantification of the gels was performed by Image J software. The mean of the wild type is normalized to 100%.Statistical analysis was calculated using student t-test, LB+22DP; **P<0.005, LB ns; nonsignificant. **B**. Conjugation efficiency of wild type and designated mutants with CRISPR-Cas targeted and non-targeted plasmids calculated as the ratio of transconjugants of targeted/non-targeted plasmids. The *P. aeruginosa* strains were grown in LB and LB+ 2,2-DP. Data shown are the mean ± SD of *n*= 3 experiments. Statistical significance was calculated using Bonferroni’s multiple comparison test (**P≤0.001).

Next, we examined the influence of PvdS on the CRISPR-Cas system in eliminating incoming nucleic acid sequences to which it has previously encountered. For this assay, we determined the efficiency of conjugal transfer of CRISPR-targeted plasmid (pUCPTSp1) and non-targeted plasmid (pUCPT) from *E. coli* S17 to *P. aeruginosa* strains. From this assay the conjugation efficiency of the targeted vs the untargeted plasmid significantly increased ∼3-fold in the *pvdS* mutants compared to the wild type under iron-depleted conditions, showing that PvdS deletion influences CRISPR-Cas targeted plasmid interference (Fig. 5B). The *pvdS* mutant exhibited a similar increase in conjugation efficiency when compared under both iron-depleted and normal conditions (∼3fold increase). However, under normal conditions, there was no difference in the conjugation efficiency between the wild type and the *pvdS* mutant. As expected, *cas3* mutation resulted in a similar effect on conjugal plasmid transfer under both iron-depleted and normal conditions (Fig. 5B). Therefore, it can be implied that PvdS regulation of the *cas* genes influences the ability of the CRISPR-Cas system to interfere with horizontal gene transfer under iron-depleted conditions.

## Discussion

*Pseudomonas aeruginosa* is characterized as a metabolically versatile bacteria capable of adapting to various biotic and abiotic stress factors in its environment by altering its regulatory systems. In this study, we highlight the ability of *P. aeruginosa* to multitask its regulatory system for adapting and thriving under iron-depleted conditions and defending against deleterious effects of horizontal gene transfer. *P aeruginosa* engages the ECF sigma factor, PvdS to regulate the CRISPR-Cas system under iron-depleted conditions (10)(11)(12). We show that iron limitation increases the *cas* gene expression (Fig. 1) as well as the CRISPR-Cas mediated plasmid interference and spacer acquisition (Fig. 5). The PvdS-CRISPR-Cas regulatory system under iron-depleted conditions, presents a well-coordinated response and an alternative defense mechanism against phage infection and HGT. This study further adds to the expanding regulatory spectrum of the CRISPR-Cas system in responding to environmental factors and the ability of bacteria to evolve defense mechanisms for survival under biotic and abiotic stress conditions.

Previous studies have shown the regulation of the CRISPR-Cas system by metabolite sensing pathways, temperature, membrane stress, oxidative stress and Quorum Sensing (23)(21)(15)(24)(25). This work further adds to previous findings indicating the induction of CRISPR-Cas expression under stress conditions compared with normal conditions (20)(23). Bacteria constantly evolve mechanisms to adapt to stress conditions in their environments, such as iron limitation. Bacteriophages are ubiquitous and their infection commonly requires prior attachment to specific receptors followed by transfer of their nucleic acids into the cell. Iron limitation induces the expression of iron transport channels (30)(31), which can serve as phage attachment sites and entry channels for nucleic acids (10)(11)(12). Hence the evolution of the regulatory system for the CRISPR-Cas system to defend bacteria under such conditions is a vital adaptive mechanism. As an adaptive immune system, expression of the CRISPR-Cas system is known to be metabolically costly to the bacteria (20)(32), Therefore, selective induction of the system under stress condition such as iron depletion, where the bacteria are most vulnerable to phage attack is more beneficial to the bacteria. As such expression of the *cas* genes is indirectly repressed by fur via the PvdS under normal conditions (Fig S3). This also reduces the risk of autoimmunity imposed by constitutive expression of the system.

In an era where increased resistance to antibiotics is on the rise, alternative therapies are being experimented for the effective treatment of pseudomonas infections. The use of phage therapy for the treatment of bacterial infections seems like a plausible venture (33). TonB-dependent siderophore uptake systems also serve as targets for antibiotics (34). The development of the trojan horse approach (9) which targets the siderophore transport channels to increase susceptibility to drugs and reduce resistance gene acquisition presents a good platform for a possible combinatory strategy with phage therapy. As seen with antibiotic therapy, bacteria continuously evolve resistance mechanisms against harmful agents. The CRISPR-Cas system remains a potent defense mechanism for bacteria and thus to develop effective phage therapy, targeting a key regulator such as PvdS may present a promising approach to downregulate the phage defense system and prevent resistance during phage therapy.

## Experimental procedures

### Strains, plasmids and media conditions

The bacterial strains, plasmids, and oligonucleotides used in this study are listed in Table S1 and S2. *P aeruginosa* UCBPP-PA14 designated in this study as PA14 was used as the wild type strain. *P. aeruginosa* strains and *E. coli* strains were grown at 37°C. Iron limited media was made by adding 300μM of the iron chelator, 2,2 -dipyridyl(2,2-DP) to the LB medium (26)(8)(35). Minimal media (MM), 0.2% (w/v) (NH_4_)_2_SO_4_, 0.41 mM MgSO_4_; 0.2% (w/v) mannitol; 40 mM K_2_HPO_4_; 14.7 mM KH_2_PO_4_, 32.9 μM FeSO_4_; 90 μM CaCl_2_; 16 μM MnCl_2_ [pH 7.2] was used for the selection of *P. aeruginosa* strains when required. When needed minimal media was supplemented with gentamicin (30μg/mL) and sucrose (10% w/v). Bacterial growth media were supplemented with antibiotics at the following concentrations: tetracycline, 10 μg/ml for *E. coli* and 50 μg/ml for *P. aeruginosa*; carbenicillin 200μg/ml for *E. coli* and 300μg/ml for *P. aeruginosa*; kanamycin, 50g/ml for *E. coli. X-gal* was added at a concentration of 50μg/ml.

The pUCPT, a derivative of pUCP19 containing *oriT* fragment from pK18, which allows for effective conjugal transfer from *E. coli* to *P. aeruginosa* was used to design the CRISPR-Cas targeted constructs. CRISPR-Cas targeted constructs pUCPTSp2n and pUCPTSp4n respectively were created by inserting the CRISPR 2 spacer1 fragments with 2 and 4 nucleotide substitutions adjacent to the GG PAM sequence, into the HindIII/EcoRI digested pUCPT. Sequences were verified by PCR and sequencing using the M13 forward and M13 reverse primers. The Mariner transposon, pBT20 which confers gentamicin resistance was used for random transposon mutagenesis to screening for regulators of the Type I-F CRISPR-Cas system. The plasmid pK18mobsacB with the gentamicin resistance cassette was used to design constructs for in-frame deletion and chromosomal *lacZ* integration.

### In-frame deletion and integrative P*cas1-lacZ* reporter construction

To create in-frame deletion mutants in *P. aeruginosa* strains, Upstream and Downstream DNA fragments flanking the region of interest in the genome were amplified by PCR using primers listed in Table S2 and ligated with the EcoRI/HindIII digested suicide vector pK18mobsacB using the MultiS cloning kit (Vazyme, China). The ligation products were transformed into *E. Coli* DH5α and positive colonies selected by blue/white screening on gentamicin (5μg/mL) and X-gal selection plates and verified by colony PCR and DNA sequencing. Selected constructs were transformed into *E. Coli* S17-1λ for conjugation with *P. aeruginosa* strains. Transconjugants were selected on Minimal medium containing gentamicin (30μg/mL), followed by the selection of in-frame deletion mutants on MM supplemented with sucrose (10% w/v). Mutants were further confirmed by PCR and DNA sequencing.

### Transposon mutagenesis

The Mariner transposon, pBT20 was transferred from *E. coli* S17 to *P. aeruginosa* PA14 (pMEP*cas1-lacZ*) strains by conjugation. Aliquots (10^−3^) of the mating mix were spread on MM agar (1.5% w/v) supplemented with Tetracycline (50 μg/ml) and 5-bromo-4-chloro-3-indoyl-D-galactopyranoside(X-gal)(50 μg/ml). Single colonies of the transconjugants were picked onto the selection plate composed of MM agar (1.5% w/v) and X-gal (50 μg/ml). Plates were incubated for 48 hours and transposon mutants visually inspected for altered blue coloration due to the up/downregulation of the *Pcas1-lacZ* activity in comparison to the wild type. Colonies with altered coloration were selected tail colony PCR and DNA sequencing using primers in Table S2. We screened over 5000 colonies on the selective media plates. Transposon insertion site mapping was achieved after BLAST search of the sequences against the *P. aeruginosa* genome (www.pseudomonas.com).

### Qrt-PCR

Cells were harvested after growth in specified media to OD_600_=1.5. RNA was extracted using the RNA extraction kit (Qiagen) according to the kit protocol. The quantity and integrity of the RNA were determined by Nanodrop and gel electrophoresis. One step Qrt-PCR reaction was performed using the Tiangen One-step SYBR Green kit (Tiangen, China) in the Applied Biosystems QuantStudio 6 Flex RT-PCR System.

### PvdS protein expression and purification

The vector pET-28b(+) (Novagen) was used for the expression of PvdS. The coding sequences of PvdS were amplified using the primer pairs listed in Table 2. The generated DNA fragments were ligated with the pET-28b(+) resulting in the fusion of the gene to the sequence encoding C-terminal His-tag. The resulting constructs, confirmed by sequencing, was transformed into *E. coli* BL21(DE3). *E. coli* BL21(DE3) colony carrying pET-PvdS was grown in LB broth supplemented with kanamycin at 37°C to OD_600_ = 0.5 and subsequently induced with isopropyl β-D-thiogalactoside (IPTG) (0.1 mM) at 16°C overnight. After overnight incubation, cells were harvested by centrifugation and bacterial pellets resuspended in cold lysis buffer (50 mM NaH_2_PO4, 300 mM NaCl, 1 mM DTT, 10 mM imidazole, pH 7.5) containing protease inhibitors (Complete mini, EDTA free, Roche) and lysed by sonification. The Cell-free supernatants were obtained by high-speed centrifugation, the supernatant was incubated with ProteinIso Ni-NTA Resin (TransGene Biotech, China) at 4 °C for 2 h. The column was washed 3-4 times with wash buffer (50 mM NaH_2_PO4, 300 mM NaCl, 1 mM DTT, 50 mM imidazole, pH 7.5) and eluted with the elution buffer (50 mM NaH_2_PO4, 300 mM NaCl, 1 mM DTT, 300 mM imidazole, pH 7.5). The protein purity was determined by SDS-PAGE analysis and dialyzed against the PBS buffer (PBS, 5% glycerol, pH 7.4) at 4 °C.

### Electromobility Shift Assay (EMSA)

The DNA fragments were generated by PCR using the indicated primers in Table S2 and end labeled with biotin using the biotin 3^I^ end DNA-labeling kit (thermo scientific) as described in the kit protocol. EMSAs were performed using the LightShift chemiluminescent EMSA kit (Thermo scientific) according to the kit protocol. Briefly, 1nm of DNA fragments were incubated with cRNAP-PvdS or PvdS and the binding buffer containing 1μg/μl Poly (dI.dC), 50% Glycerol, 1% NP-40 1M KCl 100mM MgCl_2_ and 200mM EDTA supplied with the kit for 25 minutes at 25°C. The cRNAP was added at a concentration of 1nM. The samples were then resolved on 6% native polyacrylamide gel in 0.5X TBE buffer and transferred to the nylon membrane at 380mA (∼100V) for 30 minutes. DNA was crosslinked at 120mJ/cm2 for 45-60 seconds followed by detection of the biotin-labeled DNA by chemiluminescence using the Tanon 5200 imaging system.

### Plasmid loss and spacer acquisition

The plasmids pUCPT (non-targeted naïve control), pUCPTSp2n and pUCPTSp4n (2 and 4 nucleotide substitutions in respectively) were transferred into *P. aeruginosa* strains via conjugation with *E. coli* S17. Selected colonies were cultured overnight in 5 ml LB with or without 2,2-DP and passaged for 5 days by a subculture of 20μl into 5 ml fresh media. Each passage was serially diluted and 10^−6^ dilutions plated on LB with or without carbenicillin to count colonies that retain the plasmid. *P. aeruginosa* strains initially passaged with pUCPTSp2n were further transformed with pUCPTSp4n and passaged for 5 days to assay for primed adaptation. The genomic DNA was then extracted for the identification of expanded CRISPR2 arrays using primers stated in Table S2. PCR products were resolved by 3% agarose gel electrophoresis for detection of the CRISPR array expansion.

### Conjugation efficiency assay

*E. coli* S17λ was used to transfer CRISPR-targeted pUCPTSp1 (pUCPT containing CRISPR2 spacer1) and untargeted pUCPT into *P. aeruginosa* strains. Overnight culture of *E. coli* and *P. aeruginosa* were mixed at a ratio of 1:1, washed twice and pellets resuspended in LB from which 50μl was spotted on LB agar gently to prevent splatter. The mating spot was allowed to dry and incubated for 16 hours at 37°C. The mating spot was scrapped completely and resuspended in 250 μl TSB from which serial dilutions of 10^−5^ were platted on LB and LB+2,2-DP agar with carbenicillin. The efficiency was calculated as the percentage transformants with the targeted plasmid compared with the transformants with the non-targeted plasmid.

### Beta-galactosidase assay

*P. aeruginosa* cells were grown to specified media conditions to appropriate time point and Optical density(OD_600_). Briefly, 200μl of cells were removed, pelleted and the supernatants removed completely. Subsequently, 200μl of Z-buffer (8.52g Na_2_HPO_4_; 5.5g NaH_2_PO_4_.H_2_O; 0.75g KCl and 0.246g MgSO_4_.7H_2_O, pH 7.0), 20μl 0.1% SDS and 20μl chloroform was added and vortexed for 3 min. A volume of 200μl ONPG (ortho-Nitrophenyl-β-galactoside), 4mg/ml (dissolved in Z-buffer) was added followed by incubation at 37°C for a specified time. Finally, 600μl of 1M Na_2_CO_3_ was added to stop the reaction and the absorbance measured at A_420nm_. A blank sample was composed of 200μl Z-buffer; 200μl ONPG and 600μl Na_2_CO_3._ β-galactosidase activity was calculated as (1000 x A_420_)/ (OD_600_ x Volume(ml) x time of reaction (min)) and expressed as miller units (MU).

## ACKNOWLEDGEMENT

We thank the Guangdong Province Key Laboratory of Microbial Signals and Disease Control for using their facilities to carry out the research. This work was supported by the National Natural Science Foundation of China (No. 31330002), National Key Project for Basic Research of China (973 Program, No. 2015CB150600), and Key Projects of Guangzhou Science and Technology Plan (No. 201804020066)

## CONFLICT OF INTEREST

Authors disclose no conflict of interest

